# Unraveling the temporal dependence of ecological interaction measures

**DOI:** 10.1101/2025.08.29.673018

**Authors:** Javier Aguilar, Samir Suweis, Amos Maritan, Sandro Azaele

## Abstract

Species interactions—ranging from direct predator–prey relationships to indirect effects mediated by the environment—are central to ecosystem balance and biodiversity. While empirical methods for measuring these interactions exist, their interpretability and limitations remain unclear. Here we examine the empirical matrix of pairwise interactions, a widely used tool, and analyze its temporal variability. We show that apparent fluctuations in interaction strength—and even shifts in interaction signs, often interpreted as transitions between competition and facilitation—can arise intrinsically from population dynamics with fixed ecological roles. Experimental protocols further shape these estimates: the duration of observation and the type of setup in microbial growth studies (e.g., chemostats, batch cultures, or resource conditions) systematically affect measured interactions. Considering interactions across timescales enhances interpretability: short-term measurements primarily capture direct species couplings, whereas long-term observations increasingly reflect indirect community feedback. Taken together, these results establish short-duration inferences, obtained either directly or extrapolated, as a principled way to disentangle direct from indirect interactions. Building on this insight, we propose a model-inference approach that leverages multiple short time series rather than extended longitudinal datasets.

## I. INTRODUCTION

The success of a species is rarely shaped in isolation; it emerges from a web of direct and indirect ecological interactions. Characterizing these interactions is challenging, as they involve complex relationships among multiple organisms that are shaped by a continuously changing environment^1^ and eco-evolutionary feedbacks^2^. For example, the stress gradient hypothesis suggests that as environmental stress increases, facilitative interactions become more prominent while competitive interactions diminish^3^. This implies that the environment not only determines the relative importance of species, but can also shift their roles between competitors and facilitators. Large-scale syntheses confirm broad empirical support for this prediction across taxa and ecological settings, with important implications for conservation under accelerating global change^4–7^. However, interaction shifts might result from a range of system-dependent biological mechanisms, with outcomes depending on interaction type, species traits, and environmental context^8^. The difficulty of isolating clear relationships between community members has led to theories that either ignore species interactions altogether^9–11^, or model them as random variables^12–15^. While such approaches have successfully explained some statistical patterns and broad ecological trends, constructing accurate, system-specific models—when feasible—requires robust empirical measurements of species interactions^16–19^.

Correlations or co-occurrences across species are typically the first step in inferring interaction networks^20–22^. Interaction measures are also commonly derived by fitting species abundance data with Lotka–Volterra or other population dynamics models, which are assumed to represent the underlying dynamics of the ecological communities^23–26^. Among the various model-free approaches to quantifying species interactions, estimates of the interaction matrix—defined as the effect of perturbations on a target species’ per-capita growth rate—are widely used^27–30^. Empirical estimates of interaction matrices have quantitatively supported the stress-gradient hypothesis by detecting changes in the sign of interactions, which are associated with shifts between facilitative and competitive roles^20,31–33^. However, a gap remains between the theoretical definition of interaction matrices—which assumes data of infinite resolution—and their empirical estimation, which is constrained by the finite resolution of measurements. Moreover, interaction measures do not inherently reveal whether species interactions are direct or mediated^34^; such interpretations typically rely on contextual understanding of the biological system^35^. Overall, interaction measures may be biased not only by sampling limitations, environmental effects, and experimental design, but also by the theoretical framework used to interpret them. Such biases can obscure whether the measured interactions are direct or mediated, or whether they depend on the temporal or spatial scale of observation. This leads us to ask: how do empirical conditions affect interaction measures, and what kind of information do these measures truly convey? Is it possible to determine whether interactions are direct or mediated based on the measures alone?

To address these questions, we will primarily study empirical interaction matrices extracted from synthetic data generated using paradigmatic ecological models. This approach allows us to clearly observe how interaction measures represent a perfectly known and controllable ecological process. Additionally, it enables the derivation of analytical results that provide general insights into the behavior of these interaction measures. We examine the impact of various empirical parameters, such as the resolution used to compute per-capita growth rate estimates, putting special focus on the experiment duration, i.e., the time elapsed from the ecosystem perturbation to the measurement of the per-capita growth rate. This parameter is particularly relevant in the analysis, as even if it is infinitesimal in the theoretical definition of the interaction matrix, it can be substantial in experimental setups^29,34,36^. Furthermore, it has been argued that long experimental durations reduce fluctuations in interaction estimates^34^. Through analytical calculations and simulations, we demonstrate that changes in the sign of the empirical interaction matrix can occur in purely competitive systems when the experiment duration is long enough. This is noteworthy because sign changes, traditionally linked to facilitation-competition trade-offs, may instead result from oscillations in species’ relative abundances, even while their ecological roles remain fixed.

Overall, we assess the critical role of experimental duration in characterizing ecological interactions. By temporally expanding the interaction matrix, we demonstrate that short-term measurements capture explicit couplings among ecological agents, whereas longer-term measurements increasingly reflect indirect feedback from the broader species community and environmental factors. Measuring interactions across different timescales therefore provides a more comprehensive view of system behavior: explicit species couplings can be inferred from short-term data, while indirect effects emerge over longer durations. In particular, we show that the nature of the experimental setup in microbial growth studies—such as chemostats, batch cultures, or biotic resource conditions—can significantly influence interaction estimates. Ultimately, we argue that interactions inferred at short durations, either through direct measurements or extrapolations from long-duration data, offer a systematic approach to distinguishing between direct and indirect interactions.

## II. THE EMPIRICAL INTERACTION MATRIX

In this section, we characterize the PULSE perturbation experiments and define the empirical interaction matrix derived from them^27^. Let us consider a general ecosystem consisting of *N* species with densities *x*_*i*_, *i* = 1, …, *N*. We denote by 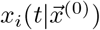 the density of species *i* at time *t* given that the state of the ecological system at time *t* = 0 was 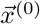. Since we aim to analyze interactions in arbitrary ecological systems, the term “species” may refer not only to organisms within a community but also to resources, either organic or inorganic. Likewise, “densities” should be understood in a general sense, representing quantities such as biomass, abundance, or other relevant measures depending on the context. This information allows measuring the percapita growth rate of species *i* at time *t*, during the time interval Δ*t*, as

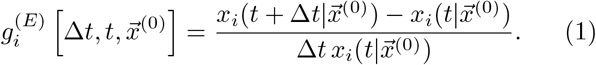

In experimental settings, the per-capita growth rate can only be estimated at a finite set of time points *t*_1_, *t*_2_, *t*_3_, … where species densities are measured. Consequently, in real applications, the parameter Δ*t* in (1) is the temporal resolution of the time series, Δ*t*_*j*_ = *t*_*j*+1_ − *t*_*j*_. PULSE experiments are based on having access to (at least) two time series of the same ecological system with different initial conditions. Specifically, the empirical interaction matrix is estimated by comparing pairs of time series, where the difference in the per-capita growth rate is evaluated between a baseline (unperturbed) evolution and a perturbed evolution. In the perturbed evolution, the initial density of species *j* is altered by an amount Δ*x*, allowing us to assess the impact of this change on the system’s dynamics,

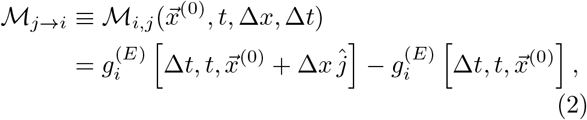

where 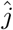 is the unit vector in the *j*-th direction. The interactions measured via Eq. (2) provide insight into the relationships between species: the sign of ℳ_*j→i*_ indicates whether changes in species *j* promote or inhibit the growth of species *i*, while its magnitude reflects the strength of this influence. However, ℳ_*j→i*_ also depends on experimental parameters that are not intrinsic to ecological dynamics. In particular, it is affected by the perturbation size Δ*x*, the total experiment duration *t*, and the sampling interval Δ*t*. When *t* is large, we will show that ℳ _*j→i*_ captures more of the system-wide feedback, incorporating indirect effects. In contrast, for small *t*, it predominantly reflects the direct impact of species *j* on species *i*, providing a clearer measure of their direct interaction, with minimal contributions from the rest of the system. In Fig. 1 we show a graphical representation of the baseline and perturbed experiments, together with experimental parameters.

**Fig. 1.**
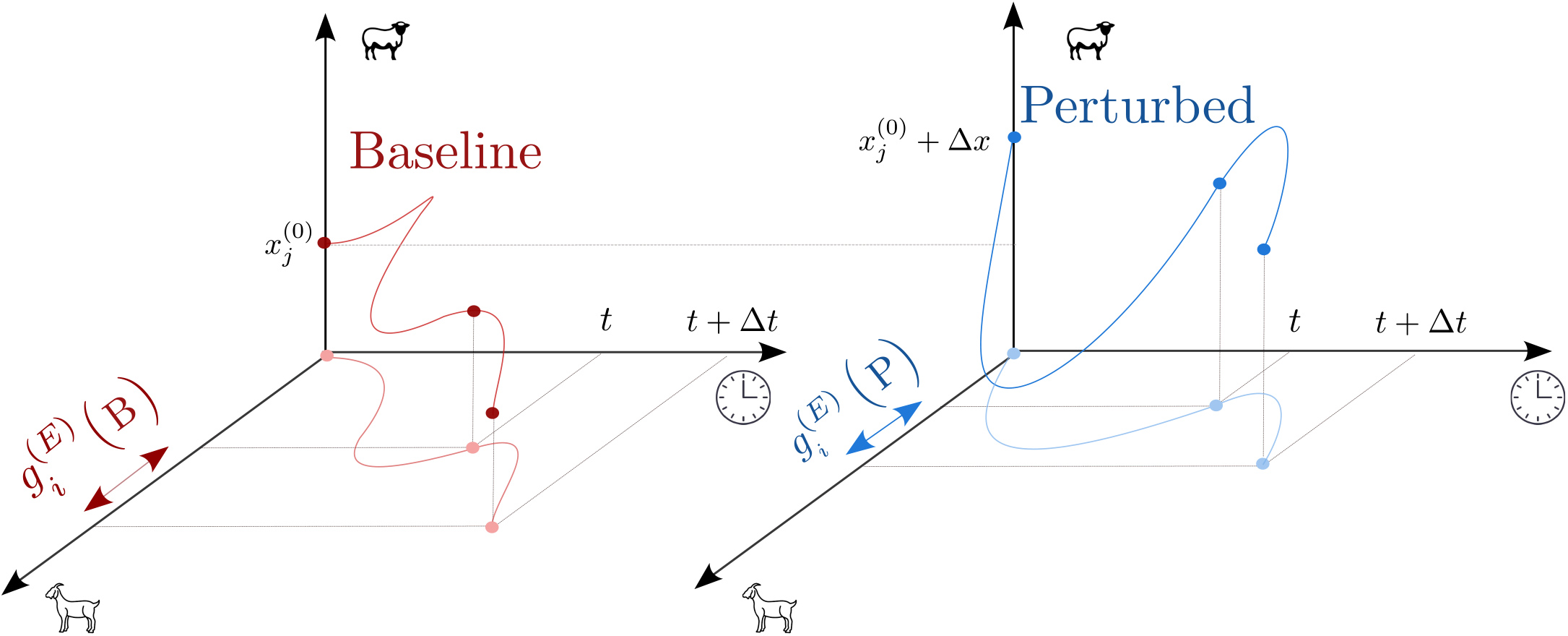
Graphical representation of the baseline and perturbed experiments. The axes represent the experiment duration (*t*), population density of the perturbed species (*x*_*j*_), and population density of target species (*x*_*i*_) to measure the per-capita growth rate 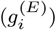.

This work focuses on understanding how experimental parameters influence the information obtained from ℳ _*j→i*_. While the interaction matrix has been studied theoretically before^27,29,30^, to the best of the authors’knowledge, little attention has been paid to the impact of these experimental parameters, which may be treated as infinitesimal in theory, but are finite in practice^27,29^.

## III. RESULTS

### III.A. Temporal variability of interaction measures

Here, we apply the interaction measure presented in Eq. (2) to two distinct ecosystems to illustrate the significance of temporal fluctuations in interactions. The first system involves the temporal dynamics of a yeast culture (S. cerevisiae) grown in a medium initially containing galactose as the sole carbon source. As the yeast consumes galactose, it produces ethanol as a byproduct, which can subsequently be used as a secondary resource^37^. This system thus represents a simple ecosystem composed of a single species interacting with two resources. We employ a consumer-resource model, fitted to experimental data from ref.^37^, to generate time series from which the interaction matrix is computed using Eqs.(1) and (2). Since we can generate time series with arbitrary resolution, we focus on the effect of the experiment duration (*t*) in the limit of small perturbation and sampling time (Δ*x*, Δ*t →* 0). Figs. 2a,b illustrate the evolution of interactions, which includes sign changes over time. Fluctuations in Fig. 2a reflect changes in the relevance of metabolic pathways. In the early stages, ethanol production exceeds consumption, resulting in a net positive interaction between ethanol and yeast. Similarly, during this initial phase, the benefits of ethanol production outweigh any competitive pressures, leading to initially positive yeast self-interactions. However, this situation changes dramatically as galactose becomes depleted. Once the system runs out of galactose, ethanol consumption surpasses production, and further increases in yeast concentration begin to inhibit both yeast growth and ethanol levels. In Fig.2b, we examine how ethanol perturbations affect yeast growth rates (ℳ _*E→Y*_). This setup compares yeast growing in a dilution containing only galactose –baseline experiment– with another evolution where a small amount of ethanol is also present –perturbed experiment–. Since resource addition typically promotes consumer growth, ℳ _*E→Y*_ is initially positive, and yeast densities are higher in solutions containing both ethanol and galactose than in those with only galactose (see Fig.2b-ii). However, the perturbed condition depletes galactose more quickly, entering faster the phase where yeast growth relies solely on ethanol (see Fig.2b-i), which supports a lower growth rate. This dynamic explains the sign shift in Fig. 2b-iii and illustrates how resource addition can eventually lead to negative interactions with consumers.

**Fig. 2.**
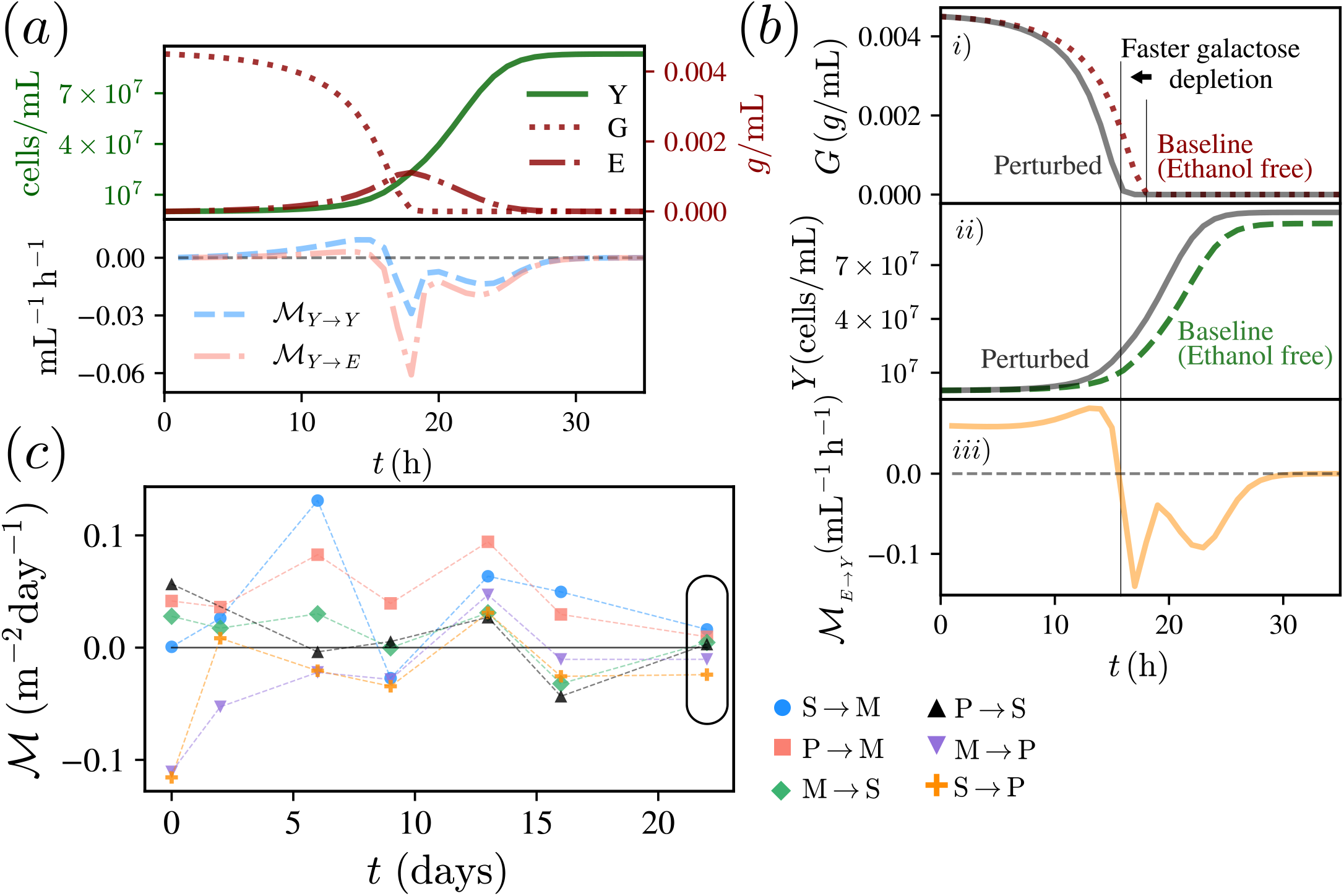
(a, top): Temporal evolution of yeast (Y), galactose (G), and ethanol (E) densities in the baseline trajectories. (a, bottom): Temporal evolution of interactions induced by yeast perturbations. (b-i): Galactose dynamics in the baseline trajectory—initially ethanol-free—and in the perturbed trajectory, which includes ethanol from the start (dotted and solid lines, respectively). (b-ii): Yeast population dynamics with and without the initial presence of ethanol. (b-iii): Evolution of the interaction measure ℳ _*E→Y*_. (c): Temporal evolution of inferred interactions among grasshoppers—<u>M. femur-rubrum</u> (M), <u>S. collare</u> (S), and <u>P. nebrascensis</u> (P). Details for parameters and model used to generate a,b in Sec. V.C.

The second dataset consists of empirical measurements of the population dynamics of three grasshopper species (<u>Melanoplus femur-rubrum</u>, <u>Spharagemon collare</u>, and <u>Phoetaliotes nebrascensis</u>) maintained in captivity. The data originate from experiments reported in ref.^38^, in which enclosures were placed in areas of natural grassland. Each enclosure contained native vegetation (serving as the grasshoppers’ shared resource) and was initialized with different population densities. These varying initial conditions allow us to infer interspecific interactions, as the grasshoppers compete for common resources. In Fig. 2c, we show that the inferred interactions oscillate over time, with many of them changing sign during the course of the experiment. Such sign changes were not reported in the original study, which only analyzed interactions at the final time point of the experiment (*t* = 28 days). That analysis was interpreted through a biological lens: dietary niche overlap among grasshoppers and possible indirect positive interactions with plants —mediated by soil enrichment— were proposed to explain the observed interaction patterns. It is noteworthy that this interpretation could be significantly affected when considering interactions at intermediate experiment durations.

Through these examples, we have examined the emergence of fluctuating interaction measures in both microscopical and macroscopical ecosystems. Understanding the meaning of interaction sign changes in the yeast–galactose–ethanol system required a great level of knowledge of organism dynamics, which was not possible to apply in the grasshopper system, which is more difficult to interpret as it is affected by noise.

These results illustrate that variability in interaction measures can arise from multiple sources in both microscopic and macroscopic ecosystems. In Fig.2a, we observe that the shifts between facilitation and competition correspond to changes in the sign of interactions, consistent with previous findings^25,32,36^. In contrast, fluctuations observed in the grasshopper system (Fig.2c) are harder to interpret. These may stem from the intrinsic randomness of the real-world data and the complex dynamics of the system as a whole. More striking is the behavior seen in Fig.2b: although resource addition typically promotes consumer growth —suggesting consistently positive interactions— we observe sign shifts that defy this expectation. In the following section, we demonstrate that such sign changes are not anomalies but a general feature, even in deterministic (error-free) models of purely *competitive* ecological systems.

### III.B. Interaction measures in synthetic datasets

In order to ensure total controllability of the ecosystem’s dynamics, we will study interaction measures in synthetic time series generated with paradigmatic ecological models. In particular, we focus on deterministic models defined through the instantaneous per-capita growth rate

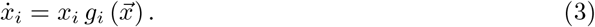

By integrating Eq. (3), we can generate time series starting from any initial condition 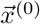 and with arbitrary sampling times. Evidently, the estimations of the per-capita growth rate (given by Eq. (1)) converge to the instantaneous growth rate that defines the model in the limit of small Δ*t*,

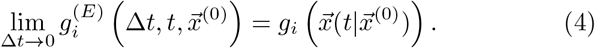

We note that Eqs. (1) and (2) make no model assumptions, since they only depend on raw data. The model in Eq. (3) just provides the time series where Eq.(2) is evaluated. This approach also allows to compute the value of ℳ_*i,j*_ analytically to gain general insights into the effects of Δ*t*, Δ*x*, and *t*. In real experiments, much care is taken to keep the sampling time and perturbation magnitude small relative to the scales of density fluctuations^30^; however, the duration of the experiment can be relatively long. Long durations of experiments have also been claimed to decrease fluctuations of interaction estimates^29^. Therefore, we perform a Taylor expansion to Eq.(2) to obtain interactions using small Δ*x* and Δ*t* while maintaining arbitrary *t* (see details in Appendix A). In doing so, we find a scalar product form of a vector only dependent on species *i* and another one only dependent on species *j*,

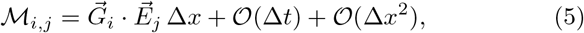

where we defined the gradient vector

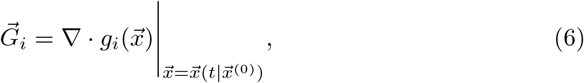

and the evolution vector

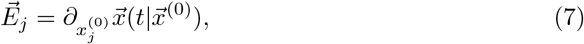

Therefore, species *j* facilitates (respectively hinders) the growth of species *i* when the evolution vector 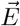 aligns with (respectively opposes) the direction of maximum variation in the growth rate of species *i* at time *t*.

Shifts in self-interactions for single-species models are straightforward to analyze, since the interaction sign coincides with the sign of the gradient vector, which reduces to a scalar in this setting (see proof in Appendix B):

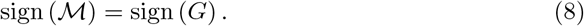

However, in multi-species models, the interaction sign depends on the evolution of both the gradient 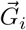, which is typically computable, and the vector 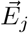, which is more difficult to evaluate analytically. This difficulty arises because computing derivatives with respect to 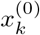 requires explicit knowledge of how the system’s trajectories depend on initial conditions. In practice, this means solving the full dynamics, which is generally infeasible for most systems. Therefore, we will approximate the vector 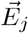 in specific time regimes to gain analytical insight.

### III.C. Oscillation of interaction measures in pure competitive systems

We focus on synthetic datasets generated with multispecies consumer–resource models without facilitation, meaning that simulated species only compete for limited resources, which themselves follow their own dynamics. Specifically, we consider a classical consumer-resource model with a general renewal function for the resources^38–41^,

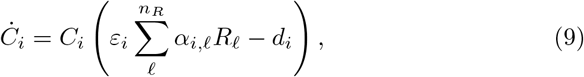

and

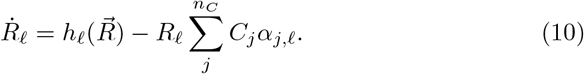

Where *n*_*R*_ and *n*_*C*_ are the number of resources and species respectively, *α*_*i,ℓ*_ is the rate of consumption of resource *ℓ* by species *i, ε*_*i*_ is the consumer’s uptake efficiency, and *d*_*i*_ is the death rate. The renewal function *h*_*ℓ*_ encodes the rate at which resource *ℓ* enters the system. All parameters and the renewal function are considered to be non-negative. Using dynamical system theory^42,43^, we demonstrate that changes in the sign of interactions among consumers can occur in systems with an arbitrary number of consumers and resources (see details in Sec. V.B). Consequently, interaction measures may switch sign even in purely competitive ecological systems. To illustrate this result, we analyze interactions in a simpler setting consisting of one consumer and one resource characterized by

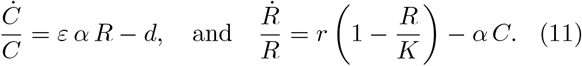

For *r >* 0, this model describes a biotic resource undergoing logistic growth. Thus, this version of the consumer–resource model can also be interpreted as a predator–prey system, where the prey (resource) is depleted by the predator (consumer). The model admits a single coexisting stationary state, 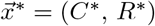. We focus on the regime where such fixed point is stable (see Appendix D), meaning that small perturbations around the stationary state decay over time, and the system returns to equilibrium. The timescales governing this relaxation are determined by two key quantities: the characteristic damping timescale

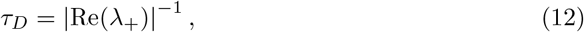

and the oscillation timescale

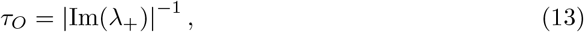

where *λ*_+_ is the eigenvalue of the Jacobian matrix with the largest real part. Notably, when

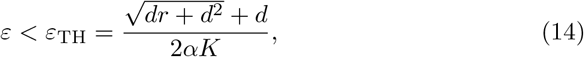

the system relaxes without oscillations 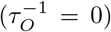, and the dynamics are dominated by exponential damping (see Fig. 3a). As shown in Appendix D, the linear stability analysis at the level of population dynamics allows us to understand quantitatively the observed variability of interaction measures.

**Fig. 3.**
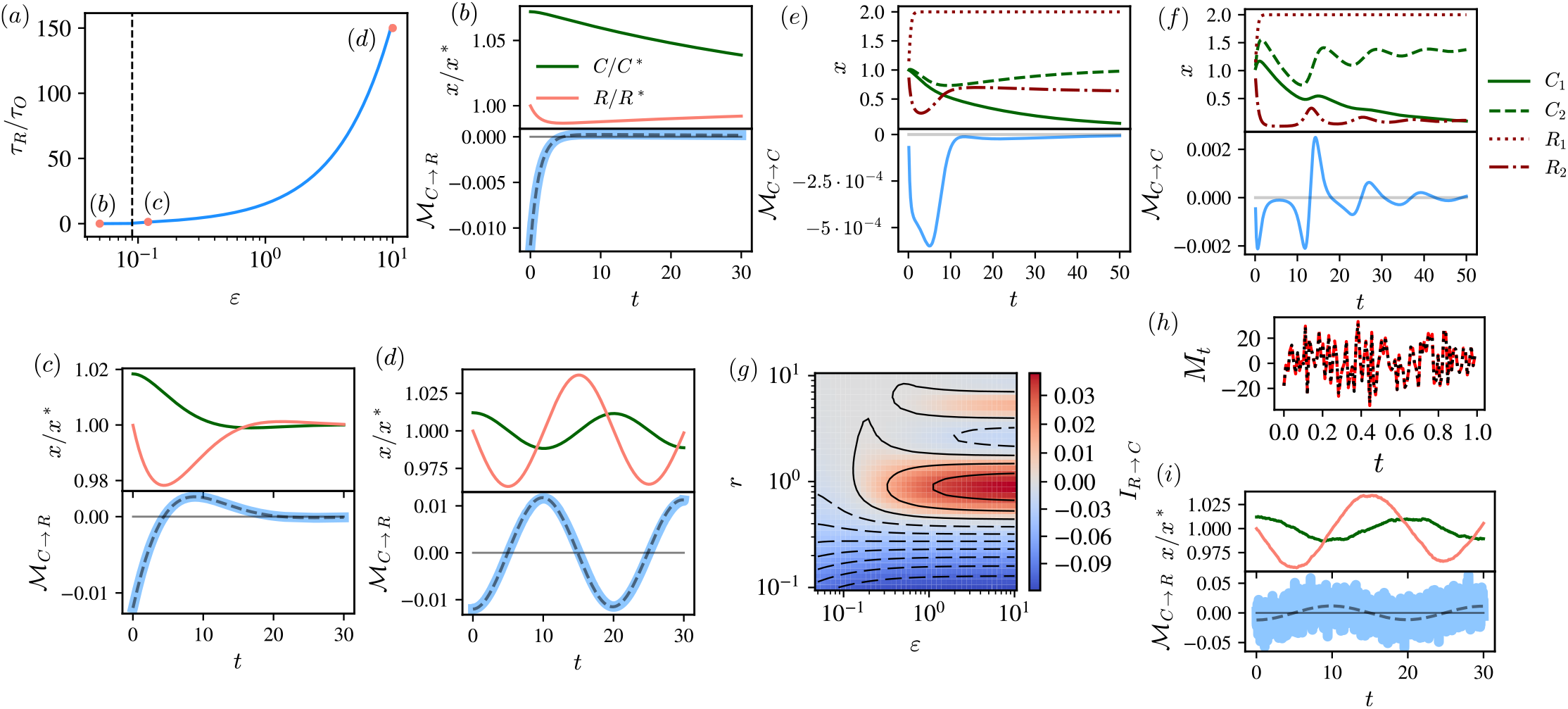
In (a), ratio between damping and oscillating time-scales in the one consumer-one resource model with biotic resource for different values of the consumer uptake efficiency. Vertical dashed line signals the threshold *ε*_*T H*_ below which there are no oscillations in the perturbation relaxation. Dots correspond to the data used in (b), (c), and (d). In (b), (c), and (d) we show instances of perturbed trajectories (top) and the associated measure of interactions (bottom). In (b) monotonic relaxation corresponds to interaction measures that do not change sign. In (c) small fluctuations rapidly damped exhibit changes of sign in the interactions. In (d), relaxation dominated by oscillations corresponds to wild oscillations in the interaction measures. Multi-species data can either show consistent interaction measures that do not change sign (e) or erratic interaction sign switches (f). In (g), integral of interactions in a time window according to Eq. (15) for different values of the consumer and resource uptake efficiency (*ε* and *r* respectively) also exhibit different signs depending on the parameters values. In (h), self-interaction measures in dataset generated with noisy process (Eq. (16)). In (i) same as in (d) but adding small noise at the population level. Dashed line in (i) signals interaction measure without noise. In the bottom plot of (b), (c), and (d) and in (h), continuous lines correspond to interactions computed from raw data, while dashed line corresponds to analytical results (Eqs. (31) and (17)). Synthetic datasets and parameter choices available in Sec. V.C.

Intuitively, the interaction ℳ _*C→R*_, i.e., how perturbations in *C* affect the growth rate of *R*, should be negative, reflecting that the addition of a consumer hinders resource growth. However, by varying the consumer’s uptake efficiency (*ε*), we observe a range of interaction scenarios — from dominantly negative (Fig.3b), to slightly oscillating (Fig.3c), and highly oscillating (Fig. 3d). This variability emerges because *ε* tunes the relative relevance of damping and oscillating time scales in the relaxation of the perturbation. Basically, when population oscillations are present, fluctuations in interaction measures reflect the inherently oscillatory nature of the population dynamics.

Overall, this simple example demonstrates that interaction measures form a dynamical system in their own right, with temporal dynamics that can be analyzed similarly to species dynamics in traditional population models. The relevance of fluctuations in these measures depends on the biological characteristics of the system — or, in the case of synthetic data, on the underlying model parameters. In multi species consumer–resource ecosystems, when there is a clear separation of timescales between consumers and resources^40^, fluctuations driven by population dynamics become negligible (see Fig. 3e). In contrast, significant fluctuations are expected when the temporal scales of consumers and resources are comparable (see Fig. 3f). In Fig. 3g, we show that the temporal integral of interactions over a finite time window [0, *t*],

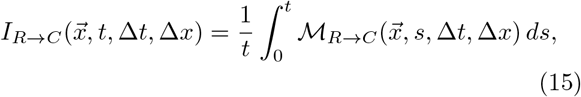

exhibits different dominant signs depending on the system’s parameters. Therefore, sign switches are not ‘averaged out’ through temporal integration; depending on these parameters, interactions can be predominantly negative or positive within a fixed time window.

So far, we have used deterministic models to evaluate ecological interactions. However, when dynamics are stochastic, the interaction matrix becomes a random variable, incorporating variability from both the process and the experimental conditions. It is still valid to apply Eqs. (1) and (2) in the stochastic setting for finite *t*, Δ*t*, and Δ*x*. As an illustration, we consider geometric Brownian motion, a minimal model of exponential growth of a small population far from saturation at its carrying capacity,

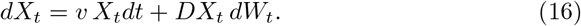

The multiplicative fluctuations in Eq.(16) prevent the population from becoming negative and are representative of environmental variability^10^. For small Δ*t*, self-interactions in this model reduce to a simple form (see Appendix G),

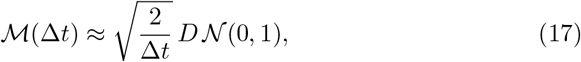

where 𝒩 (*µ, σ*^2^) is a normal random variable with mean *µ* and variance *σ*^2^. Therefore, the self-interaction takes both positive and negative values erratically (Fig. 3h). We note that the infinitesimal limit in Eq.(4) may not exist in time series with abrupt fluctuations. For example, time series generated by white-noise-driven paths such as Eq.(16) do not have well-behaved infinitesimal growth rates or infinitesimal self-interactions, as evidenced by the divergence of Eq.(17) in the limit Δ*t* → 0. However, well-behaved empirical growth rate estimates over stochastic paths may still occur, as in synthetic time series generated with stochastic processes with correlated noise — a more natural choice for ecological modeling, as discussed in AppendixH.

In summary, random fluctuations in population dynamics can cause sign changes in interactions that are unrelated to facilitation–competition trade-offs. Indeed, in Fig.3-i we show that small fluctuations in a simple consumer–resource model considerably increase temporal variability, an effect that can be even more pronounced if stochastic amplification plays a role^44^. Averaging over multiple realizations reduces this variability:

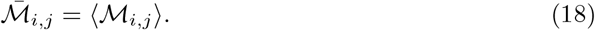

This average interaction matrix can be interpreted as a statistical correlation,

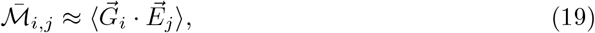

highlighting that 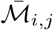 expresses a statistical — not necessarily causal — relationship between the growth rate of species *i* and *j*.

### III.D. Interaction measure design

In the previous section, we demonstrated that sign changes in interaction measures can arise from intrinsic oscillations during relaxation processes. This presents challenges for interpreting the outcomes of PULSE perturbative experiments, as identical species and environmental conditions can lead to significantly different results depending on the experiment’s duration. Consequently, a natural question arises: what experiment duration should practitioners use to ensure that interaction measures are informative? In the following, we show that by performing interaction measurements at multiple experiment durations, one can obtain a more comprehensive understanding of system dynamics, capturing not only species-specific behavior but also their interactions with the environment and the experimental setup. To do so, we turn our attention to interactions occurring over short experiment durations. This analysis is carried out through the expansion of Eq. (5) as a power series in the experiment duration, *t*, and in the limit of vanishing Δ*t*,

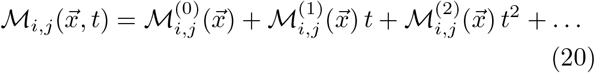

The zeroth-order in the expansion simply becomes the derivative of the per-capita growth rate evaluated at the perturbation point (see Appendix E for details):

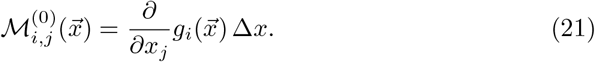

Therefore, the zeroth-order term in the expansion depends solely on the gradient vector 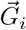 and captures the explicit couplings between species *i* and *j* present in the model used to generate the synthetic data. As such, this term represents the direct interactions between species. In contrast, the higher order terms in time in Eq. (20) depend on both 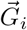 and 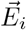, and they encode the effects of indirect interactions among species, environmental dynamics, and experimental protocols. Specifically, if there is no explicit coupling between species *i* and *j*, i.e., if 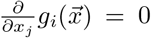, then the leading term in the expansion becomes (see Appendix E):

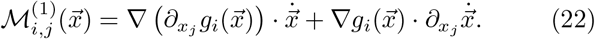

For a broad class of systems, Eq. (22) simplifies, and 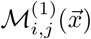 coincides with the instantaneous interaction matrix defined in Ref.^36^, which characterizes indirect interactions among species mediated by the environment (see Appendix E.1).

The above analysis provides a method to determine whether an interaction measure reflects direct or indirect interactions between species. The key idea is to exploit the fact that these two types of interactions behave differently in the limit *t*→ 0. While indirect interactions should tend to zero near *t* ≈ 0, direct interactions should intercept the y-axis at a finite value corresponding to 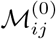 (see sketch in Fig. 4a). In Fig. 4b, we show the different behavior of direct and indirect interactions using a multi-species consumer resource model. The interactions ℳ _*R→C*_—which quantify how resource perturbations affect consumer growth—tend toward a positive value as *t →* 0, reflecting the explicit coupling between consumers and resources. In contrast, the interactions ℳ _*C→C*_ tend towards zero, indicating that the consumer-consumer interactions are indirect and mediated by resource availability. In Fig. 4c, we show the case where ℳ _*C→C*_ interactions are only available at finite experimental times. Although measures with finite *t* are always finite and negative, reflecting mediated competition, the uncoupled nature of consumers is revealed by the limit *t* → 0, which is accessed through extrapolation.

**Fig. 4.**
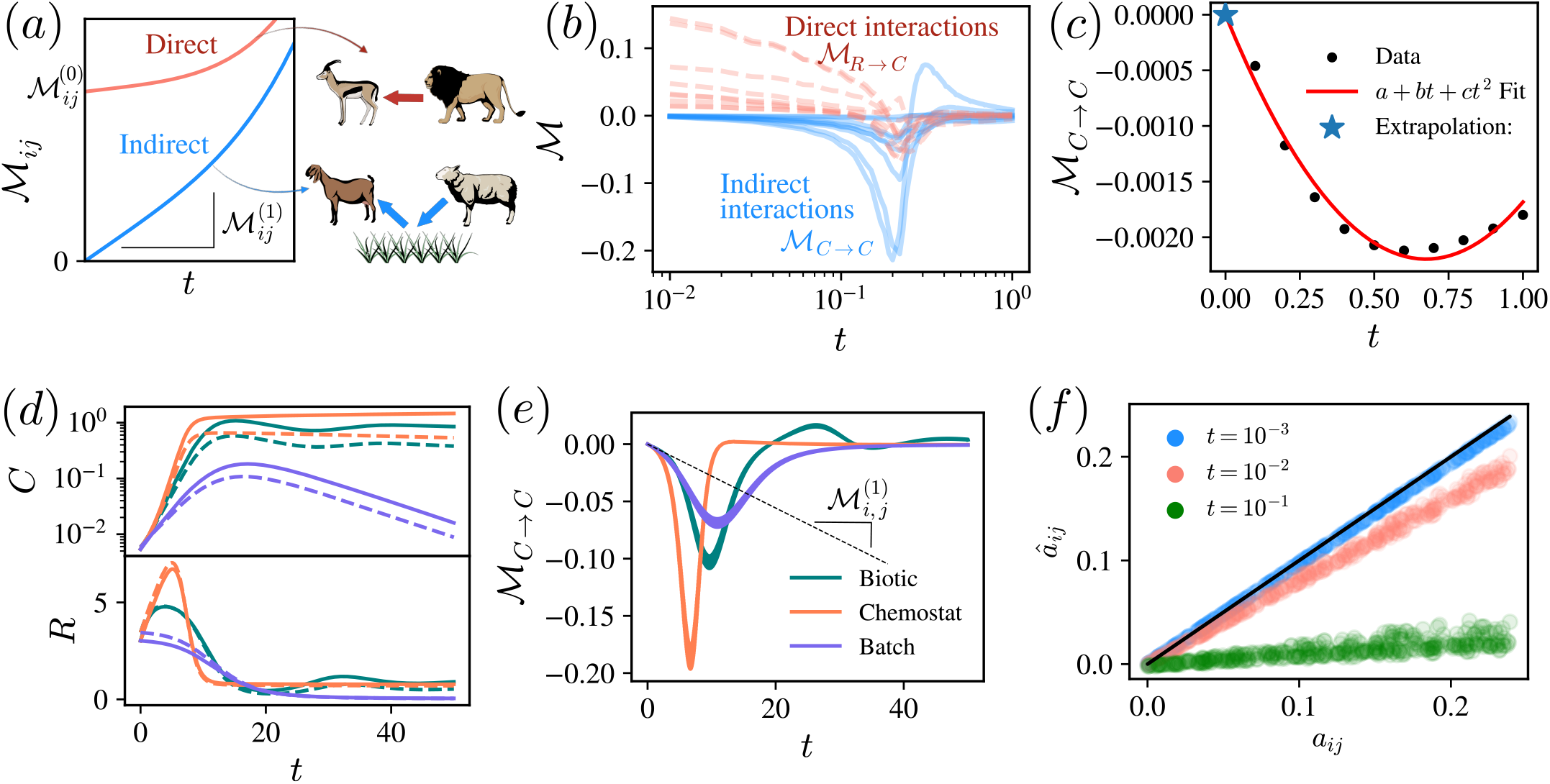
In a, sketch illustrating the hallmarks of direct and indirect interactions. In the limit *t →* 0, direct interactions approach a finite value 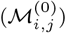, while indirect interactions vanish proportionally to 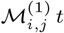. In b, interaction measures for short experiment durations in a multi-species consumer-resource model. The interactions ℳ_*C→C*_ tend to zero, while ℳ_*R→C*_ converge to a positive value as *t →* 0. In c, interaction measures between consumers (*n*_*R*_ = *n*_*C*_ = 2) at various discrete experiment durations (dots); data is fitted with a parabolic function (solid line), and extrapolation to *t* = 0 correctly recovers the interaction limit 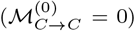. In d, baseline trajectories in a two-consumer, two-resource model, with different colors representing resource renewal types (biotic, chemostat, batch), while coupling constants are identical across all types. In e, interaction measures across renewal types corresponding to trajectories in d. In f, inference of coupling constants in a 20-consumer, 20-resource model using interaction measures with three different experiment durations; the x-axis shows true parameter values, the y-axis shows inferred values, and the solid line indicates perfect inference. Synthetic datasets and parameter choices available in Sec. V.C.

In experimental realizations of consumer–resource communities—both those involving macroscopic organisms and microbial communities—the experimental protocol is often largely defined by the nature of resource input into the system. One approach is to sample species’ growth rates under natural resource conditions (see e.g.^34,45^). To achieve more controlled conditions, experimentalists can instead isolate communities from their natural environments and (i) study population decay when only an initial quantity of resources is provided without external input^19,38,46,47^, or (ii) provide a continuous input of selected resources to investigate stationary properties^24,25,48^. Since we have shown that population dynamics influence the outcomes of interaction measures, we expect that the choice of experimental protocol may significantly affect these measures. In the consumer-resource model defined in Eqs (9) and (10), different experimental protocols can be analyzed through the choice of the resource renewal function *h*_*ℓ*_. In particular, we focus on three types of resource renewal that can be found in experiments of microbial communities:

- **Batch culture:** There is no renewal of resources. An initial quantity of resources is present and decays due to consumption. Thus,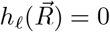.
- **Chemostat:** Abiotic resources are introduced at a constant rate, establishing a steady state. This corresponds to a constant renewal function, 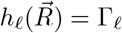
- **Biotic resource:** Resources follow their own natural dynamics and coevolve with the consumers. We model this with logistic growth,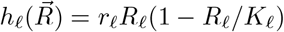.

By computing the explicit parameter dependence of the expansion in Eq. (20), we find that the first two terms behave identically across all experimental protocols, as they do not depend on the resource renewal functions or on the consumers’ death rates (see Appendix E.2):

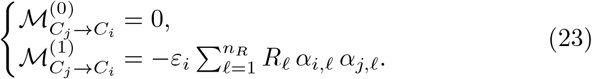

In Eq. (23), ℳ ^(0)^ reflects the absence of direct interactions among consumers, while ℳ ^(1)^ captures the expected negative indirect interactions mediated by resource competition. Notably, ℳ ^(1)^ generalizes the interaction structure found in the Lotka–Volterra approximation of the consumer–resource model with fast resource dynamics^40^. In Appendix E.2 we show that the next term in the expansion, 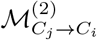, already depends on the renewal function and can be either positive and negative. In Figs. 4 d,e, we show trajectories and interactions in a two-consumer, two-resource model with different resource renewal functions but identical consumer-resource coupling constants, thus representing constant consumer-resource relationships under different experimental conditions. Interaction measures in all experimental settings behave similarly at very short times, when consumer-resource coupling dominates, in line with Eq. (23). However, as the experiment progresses, the impact of the experimental protocol on interaction measures grows. Additionally, we show that changes of sign in the interaction could critically depend on the renewal function, suggesting that sign changes could be artifacts induced by the experimental setting.

### III.E. Model inference from interaction measures

So far, we have demonstrated how empirical estimates of the interaction matrix behave on synthetic data generated by models. However, we could also ask which types of model can be inferred from measurements of the interaction matrix. Using the normalized version of the interaction matrix^30^,

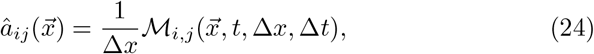

then the zeroth order model inferred from interactions is

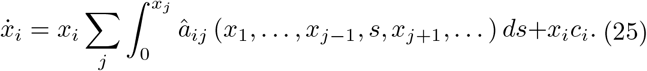

The derivation of Eq.(25), detailed in Appendix F, relies on the fact that in the limit *t* →0, the relationship between interaction measures and per-capita growth rates becomes trivial. This inference method cannot capture one-body processes (such as death, asexual reproduction, or migration), which remain undetermined and are parameterized by the constants 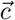. The advantage is that these constants can be estimated from single-species experiments. In Fig. 4f, we show that Eq. (25) can be used to infer the coupling parameters (*α*’s in Eq. (9)) in a consumer-resource model with many species. The errors of the parameter estimation depend on the experiment duration *t*.

We note that the model inferred from Eq. (25) is quite general and can, in principle, account for multi-body interactions. This generality arises because the estimated interaction coefficients *â*_*ij*_ may depend on the densities of all species, denoted by 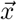. However, the method recovers the classical Generalized Lotka-Volterra model when the coefficients *â*_*ij*_ are constant:

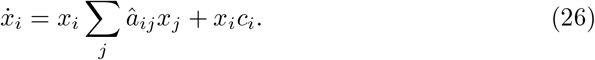

Equation (26) illustrates the universal emergence of Generalized Lotka–Volterra dynamics from empirical measurements of pairwise interactions, provided these interactions are not sampled across different initial conditions. In other words, when the estimators *â*_*ij*_ do not depend on species densities—either because densities are sufficiently small for independence to hold approximately, or because interactions are measured only from baseline trajectories with constant initial conditions—the inferred model necessarily reduces to a Generalized Lotka–Volterra form.

## IV. DISCUSSION

Measurements of the empirical interaction matrix are influenced by the experimental protocol, particularly the experiment’s duration. In short-duration experiments, the interaction matrix directly captures the causal relationships between ecosystem components. However, as the experiment duration increases, these measurements increasingly reflect correlations between per-capita growth rates and density fluctuations, rather than direct causal interactions. This shift does not mean that longer-duration measurements are uninformative. Instead, the emergence of correlations allows the analysis of indirect interactions that may not be evident in direct causal relationships between species^49^.

That said, these measurements should be interpreted with caution. For instance, changes in the sign of interactions might be seen as support for the stress-gradient hypothesis, which posits that interactions are a mix of facilitation and competition, with the balance shifting based on environmental context. However, our findings suggest that even when species roles are fixed—meaning there is no explicit dependence on environmental conditions in growth rates and no facilitation-competition trade-offs—the sign of interactions can still change due to intrinsic fluctuations in the system’s dynamics. This finding emphasizes that ecologists should consider the possibility of intrinsic temporal fluctuations when interpreting changes in interaction signs observed in empirical data. Moreover, we have shown that measures of interactions can be influenced by experimental decisions, such as how nutrients are supplied to species in controlled growth experiments. Thus, clearly documenting and standardizing experimental protocols is crucial to ensure comparability and reliability of inferred ecological interactions across different studies.

Identifying whether an interaction measure captures direct or indirect species interactions is a challenging task that has traditionally relied on a detailed understanding of the underlying biological processes. Based on such understanding, researchers infer how to interpret the resulting interaction measures^34,45,50,51^. Here, we propose a systematic approach to distinguish the nature of interactions. By evaluating interaction measures at finite times, we can extrapolate their behavior in the short-time limit, which provides direct insight into the directness of the interactions. Practically, this approach offers researchers a robust analytical method for interpreting ecological experiments, even when direct short-term measurements are challenging or noisy.

The interaction matrix does not assume any specific underlying model for its empirical estimation. This raises the natural question of whether such measures can provide insights into plausible generating models for the data, which we address with a proposed inference method. The method is based on sampling short time series over many initial conditions that are then perturbed. This approach offers a paradigm shift compared to typical inference methods, which often require rich longitudinal data—i.e., long and stationary time series. A key feature of the proposed method is that its information is additive: it disentangles single-species and multi-species dynamics, making it possible to separate data from individual species and multi-species experiments, and then combine the information. However, it is important to acknowledge that the accuracy of inferred models is sensitive to the density and precision of data collected at very short timescales, which may be challenging to achieve in field studies.

The main takeaway from this work is the idea that the experimental protocol is intrinsically linked to the outcomes of measurements, even when assuming error-free data. Therefore, parameters characterizing the experimental setting are just as crucial as those governing species behavior in determining the dynamics of measured interactions. We have shown that mathematical models not only aid in understanding the general properties and underlying mechanisms of ecological systems, but also provide a framework to assess the extent of information in empirical measurements. Ultimately, integrating models with empirical estimates offers a valuable perspective for designing more informative experimental protocols and improving the quality of empirical data.

## V. METHODS

### V.A. Data and code availability statements

Code used to generate synthetic data together with the particular datasets used in figures is available in^52^. Grasshopper’s dataset is also present in^52^, but it was extracted from ref.^38^.

### V.B. Interactions in consumer resource model

To compute interactions in the general consumer resource model defined in Eqs. (9) and (10), we can use the framework developed in section III.B simply considering a system with *N* = *n*_*C*_ + *n*_*R*_ entities with densities

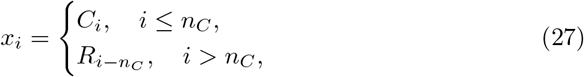

and per-capita growth rates

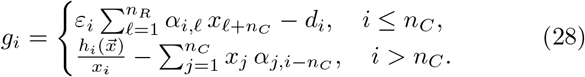

An interesting feature of this sort of consumer-resource models is that the gradient vector associated with consumers is time-independent:

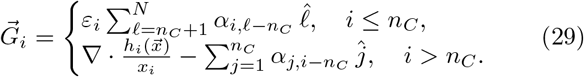

Therefore, any temporal variability in interaction measures associated with consumer growth is driven by the evolution vector 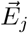. The behavior of the evolution vector becomes more transparent at the stationary regime, where it can be approximated through linear response theory. Assuming initial conditions in the vicinity of a fixed point 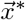 such that

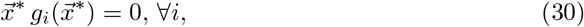

the interaction matrix in the small Δ*t* and Δ*x* limit becomes

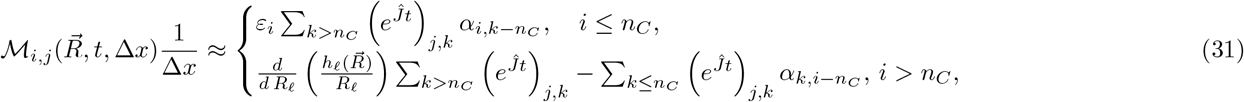

where *ĵ* is the Jacobian,

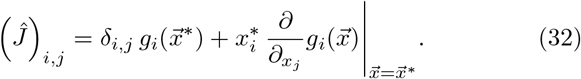

Eq. (31) was obtained linearizing the dynamics around 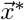, as shown in Appendix C. The elements of the exponential matrix appearing in Eq. (31) can be either positive or negative in multi-species models^53,54^. Therefore, we expect to observe changes in the sign of interactions measures even in the case of purely competitive ecosystems.

### V.C. Parameters used in figures

- <u>Fig.2 a,b</u>: Data generated with yeast-galactose-ethanol model of ref.^37^,

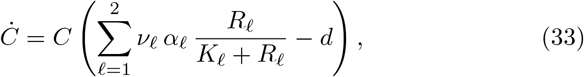

and

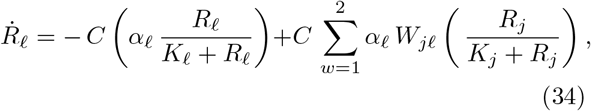 Where *C* represents the density of yeast, values and parameters with *ℓ* = 1 are associated with galactose, and those with *ℓ* = 2 with ethanol. Parameter’s values correspond to the fit to real data also performed in ref.^37^. *d* = 10^*−*5^ (hours^*−*1^), *K*_1_ = 6.73 ·10^*−*4^ (g mL^*−*1^), *K*_1_ = 10^*−*3^ (g mL^*−*1^), *ν*_1_ = 2.03·10^9^ (g^*−*1^), *ν*_1_ = 4.35 ·10^10^ (g^*−*1^), *α*_1_ = 1.13 ·10^*−*10^ (g hour^*−*1^), *α*_2_ = 1.21· 10^*−*11^ (g hour^*−*1^), *W*_1,1_ = *W*_2,2_ = *W*_2,1_ = 0, *W*_1,2_ = 0.3, *C* (*t* = 0) = 162734.08 (m^*−*2^),*R*_1_ (*t* = 0) = 4.5 · 10^*−*3^ (g mL^*−*1^), *R*_2_ (*t* = 0) = 0. Regarding the sampling times and perturbations, Δ*t* = 1 (hour), Δ*x*_*R*_ = 10^*−*4^ (g mL^*−*1^), Δ*x*_*C*_ = 10^4^ (m^*−*2^).
- <u>Fig.3</u>: Data generated using model in Eqs. (9) and (11). In a-d, *n*_*C*_ = 1, *n*_*R*_ = 1, *a* = 1.2, *K* = 2, *d* = 0.1, *r* = 1. In b,c, and d, respectively, *ε* = 5 ·10^*−*2^, *ε* = 0.12, and *ε* = 10. Initial conditions coincide with stable fixed points (Eq. (D3)) In e-f, *n*_*C*_ = 2, *n*_*R*_ = 2, *α*_1,1_ = 0.05, *α*_1,2_ = 1, *α*_2,2_ = 2, *α*_2,1_ = 0, *K* = 2, *d* = 0.1, *r* = 3, *R*(*t* = 0) = *C*(*t* = 0) = 1, *ε* = 0.08 in e, and *ε* = 0.5 in f. In g, same parameters as a-d. In h, data generated with Eq. (16) with *v* = 1, *D* = 1, *X*_0_ = 1. In i, same as in d with additive noise, diffusion constant *D* = 10^*−*4^.
- <u>Fig.4</u>: Data generated using model in Eqs. (9) and (10). In b, *n*_*R*_ = *n*_*C*_ = 10, renewal function 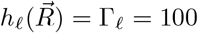. Defining *U*_[*a,b*]_ as an uniformly distributed random variable in the interval [a,b], 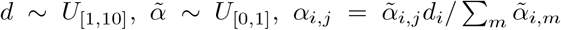 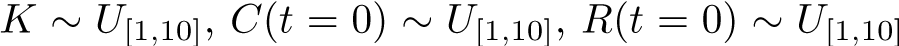. In c, same as in Fig.3f. In 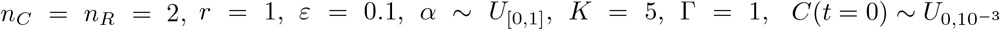. In f, *n*_*C*_ = *n*_*R*_ = 20, *α ∼ U*_[0,2]_.

## VI. ACKNOWLEDGMENTS

We thank Sara Mitri, Davide Bernardi and Luca Martinoia for their useful comments. J.A., S.A. and A.M. acknowledge financial support under the National Recovery and Resilience Plan (NRRP), Mission 4, Component 2, CUP 2022WPHMXK, Investment 1.1, funded by the European Union – NextGenerationEU – Project Title: “Emergent Dynamical Patterns of Disordered Systems with Applications to Natural Communities”. S.S. acknowledges financial support under the National Recovery and Resilience Plan (NRRP), Mission 4, Component 2, Investment 1.1, Call for tender No. 104 published on 2.2.2022 by the Italian Ministry of University and Research (MUR), funded by the European Union – NextGenerationEU – Project Title: Anchialos: diversity, function, and resilience of Italian coastal aquifers upon global climatic changes – CUP C53D23003420001 Grant Assignment Decree n. 1015 adopted on 07/07/2023 by the Italian Ministry of Ministry of University and Research (MUR).

## VII. AUTHOR CONTRIBUTIONS

JA, SS, AM, and SA conceived and designed the study. JA performed the calculations, simulations, and data analysis. JA, SS, AM, and SA wrote the manuscript, and all authors reviewed and approved the final version.

## Appendix A: Analytical expression for interactions evaluated on synthetic data

In this section, we provide the proof for Eq. (5) from the main text, which enables the evaluation of the empirical interaction matrix when applied to a synthetic dataset generated by a model with a known per-capita growth rate. The derivation makes use of multidimensional Taylor expansions. Consider that the per-capita growth rate is evaluated on a vectorial function 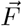 that depends itself on a vectorial quantity 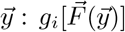. Then, let us perturb the *j*^*th*^ component of *y* in an small amount Δ*y*,

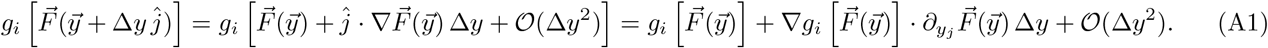

Therefore,

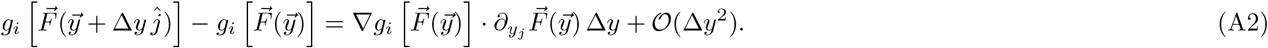

The above expression directly leads to Eq. (5) by substituting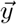 by 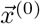, and 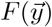 by 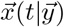.

## Appendix B: Interactions in single species systems: self interactions

As discussed in section III.B of the main text, it is not generally possible to compute the value of empirical interactions measured over synthetic data as this entails solving the entire dynamics of the species densities. However, this is feasible in one dimensional systems, where one would obtain *x*(*t*|*x*^(0)^) solving the equation

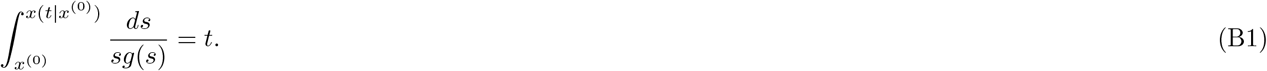

Moreover, it is possible to extract information about interaction measures even without solving Eq. (B1). Let us re-write eq. (3) in a one-dimensional setting and making explicit the dependence on the initial condition

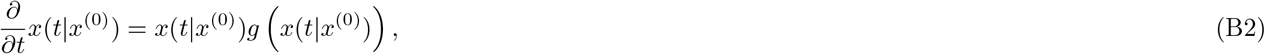

deriving with respect to *x*^(0)^ the above equation and assuming that derivatives can be exchanged, we can write an ordinary differential equation for 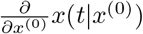,

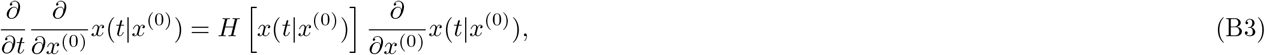

with

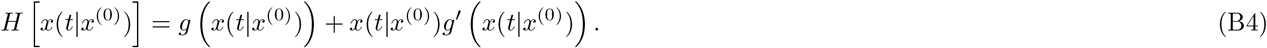

Therefore,

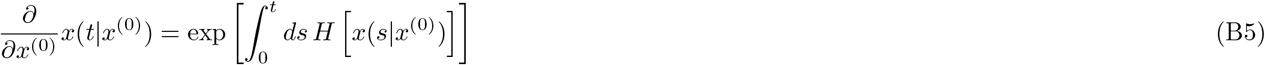

where we fixed the integrating constant imposing 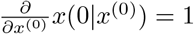. Using this result we can rewrite eq. (5) as

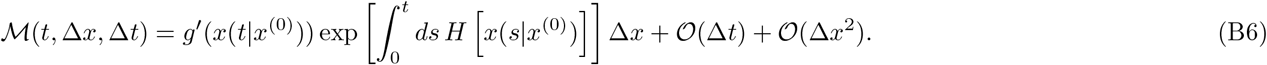

Since the range of *H*(·) belongs to the reals, the exponential in the above equation is always positive and changes in the sign of the interactions only depend on the sign of the derivative of the per-capita growing rate. Therefore, in this setting changes of sign in the self-interactions only can arise from competition-facilitation trade-offs.

### B.1. Interactions in single species: logistic growth

To illustrate measures of self-interactions in single-species growth, we analyze the logistic growth:

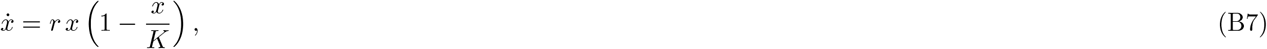

with the solution

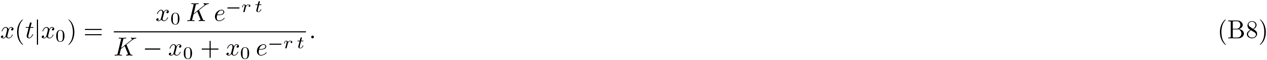

Eq. (B8) can be used to generate time series with the desired sampling time Δ*t*, magnitude of perturbation Δ*x*, and experiment duration *t*. Moreover, having the explicit solution allows us to evaluate Eq. (5) exactly. In the case of single-species evolution, the interaction matrix reduces to a single value that accounts for self-interactions — i.e., the effect of small perturbations in the species population on the overall density growth.

In Fig. 5 we show that Eq. (5) properly describe the self-interactions measured from realizations of the process. The sign of self-interactions is always negative, being consistent with the system’s dynamics, as increasing the density only decelerates the per-capita growth if *x*^(0)^ *< K* or accelerates the per-capita decay if *x*^(0)^ *> K*. We also observe that interactions converge to zero as the experiment duration *t* grows, this trend is expected as both the baseline and perturbed trajectories will end up at the same stable fixed point *x*^*∗*^ = *K*. Indeed, in the logistic growth *g*^*′*^(*x*) = − *r/K*, and thus changes of sign of self-interactions are not possible.

**Fig. 5.**
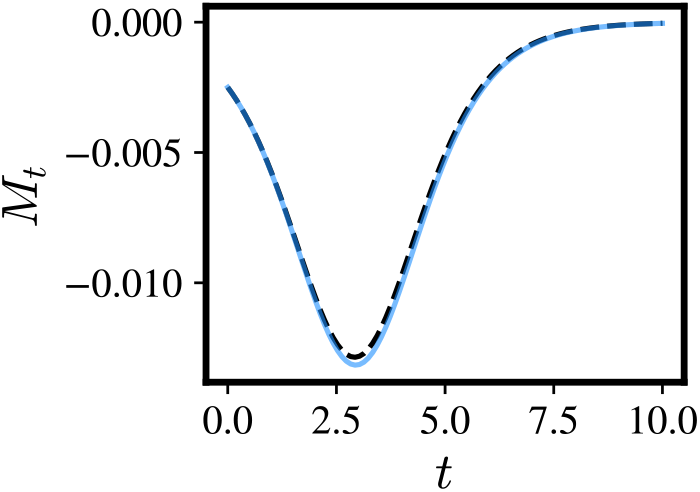
Self-interactions in the logistic model. Interactions computed on synthetic data generated with Eq. (B8) (continuous blue lines) and using the analytical approximations results (dark dashed lines) from eq. (5).

## Appendix C: Interaction measures close to a fixed point

In this section, we consider initial conditions in the vicinity of a fixed point 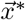 such that

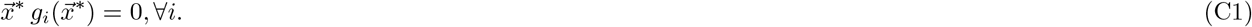

When 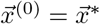, the equation of motion for the evolution of the perturbed time series can be linearized. Considering

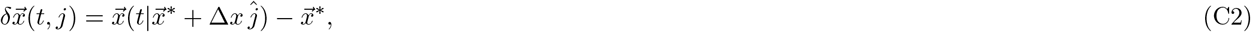

then

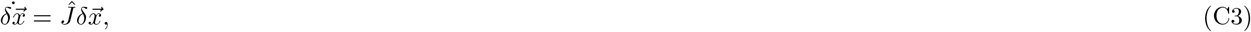

where *ĵ* is the Jacobian,

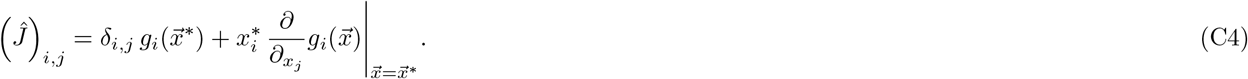

Therefore, trajectories in the vicinity of fixed points looks like

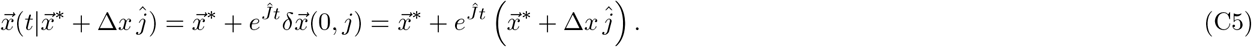

The derivative of the above expression with respect to Δ*x* gives the Evolution vector, which we can identify as the *j−*th column of the exponential matrix 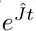,

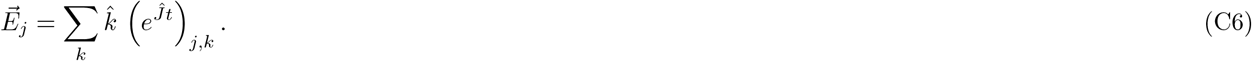

Therefore, interactions in the vicinity of fixed points can be computed as

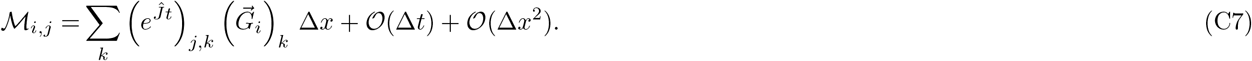

## Appendix D: Linear analysis in one consumer-one resource model

In this section, we make the linear stability analysis for the simple model

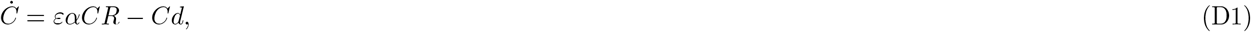

and

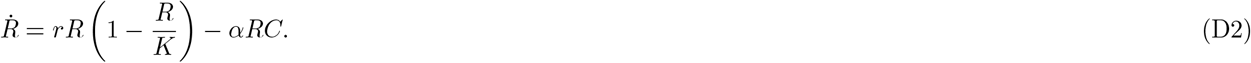

Among the model’s five parameters—*r, K, α, d*, and *ε*—three can be eliminated through rescaling of the variables *R, C*, and *t*. This leaves only two independent parameters that govern the possible behaviors of the interactions. For our analysis, we choose these to be *r* and, in particular, *ε*.

The eigenvalues of the Jacobian of the dynamics evaluated at the coexistence fixed point,

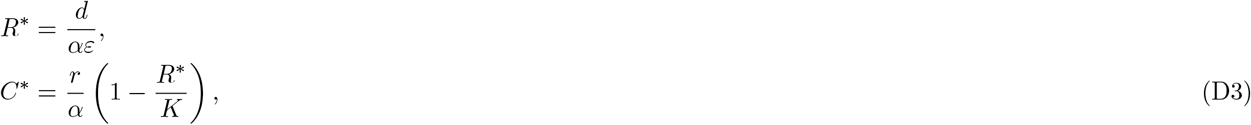

reads

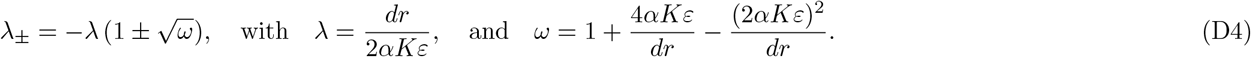

The coexistence fixed point is always stable, as the real parts of both eigenvalues are negative. However, this fixed point has positive components only if *R*^*∗*^ *< K*, or equivalently, if 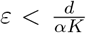. Therefore, we focus on this parameter region.

The time scales of damping and oscillations characterizing the relaxation towards equilibrium are related with the real and imaginary parts of the eigenvalues respectively

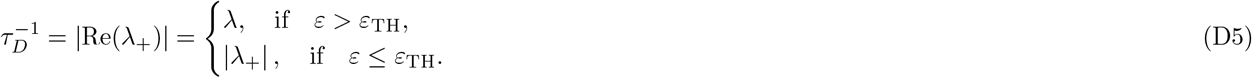

and the oscillation timescale,

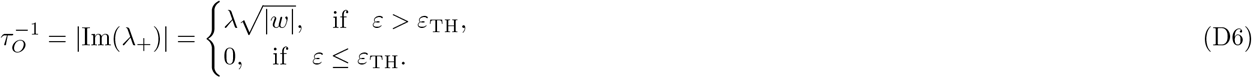

with 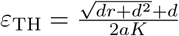.

*τ*_*D*_ characterizes the exponential decay toward the coexistence fixed point, with larger *τ*_*D*_ indicating a slower decay. Similarly, *τ*_*O*_ determines the frequency of oscillations, with larger *τ*_*O*_ corresponding to greater temporal distance between consecutive peaks. The ratio

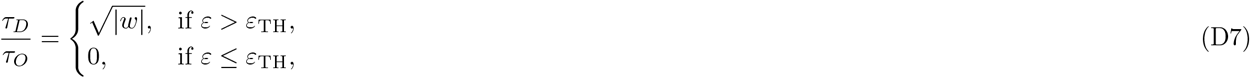

describes the relative importance of damping versus oscillatory behavior. A small value of *τ*_*D*_*/τ*_*O*_ indicates that oscillations are slower than damping, and thus perturbation relaxation is expected to be mostly monotonic. In contrast, a large *τ*_*D*_*/τ*_*O*_ corresponds to a highly fluctuating relaxation, as illustrated in Fig. 3.

The oscillation threshold *ε*_TH_ increases with *r*. Therefore, as *r* increases and *ε* decreases, oscillations become less significant. In this limit, the resource evolves much faster than the consumer, allowing the consumer dynamics to be effectively decoupled from the resource. The resulting approximation is:

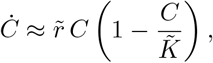

where 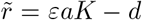, and 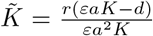. Therefore, in this time-scale separation limit, oscillations are suppressed as the system effectively lives in a one-dimensional space.

## Appendix E: Expansion of the empirical interaction matrix for short experiment durations

In this section, we provide more details about the derivation of the equations apearing in section III.D. Integrating the model definition we obtain,

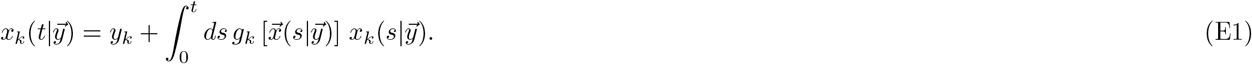

Where 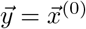.

The Taylor series expansion to first order reads

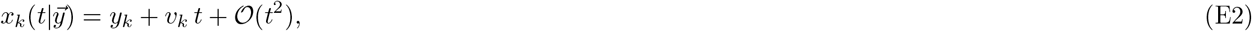

were we defined the ‘velocity’

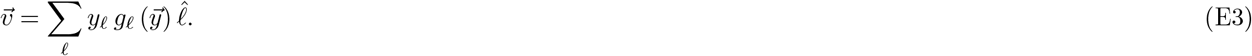

Inserting the above series-expansion again in Eq. (E1), we increase one order in the expansion

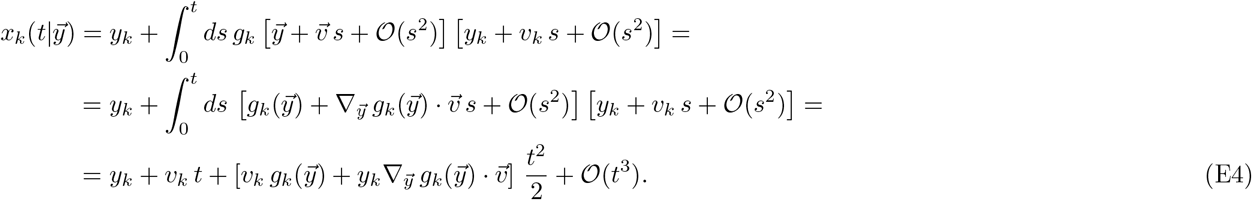

Using

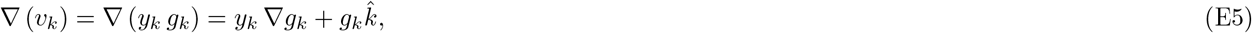

then

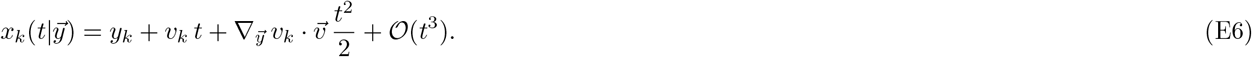

which leads to the following expression for the components of the evolution vector:

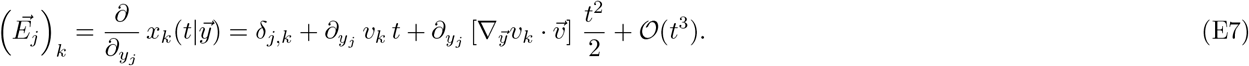

Next, using eq. (E6) we approximate the gradient to second order:

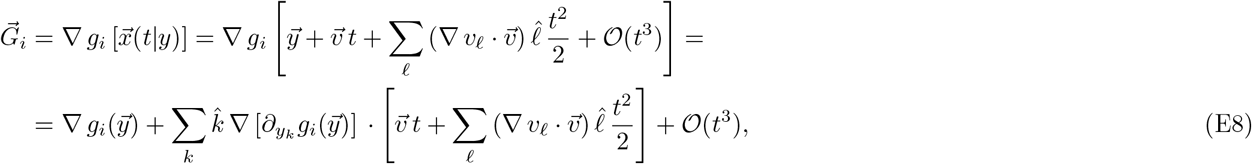

Inserting Eqs. (E7) and (E8) in Eq. (5) we can get the coefficients of the interaction matrix,

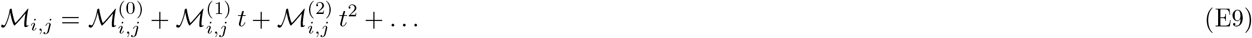

In particular,

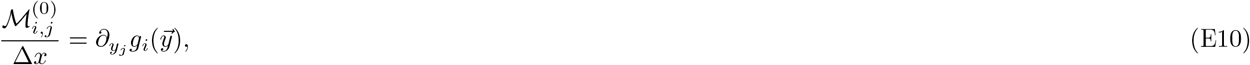

and

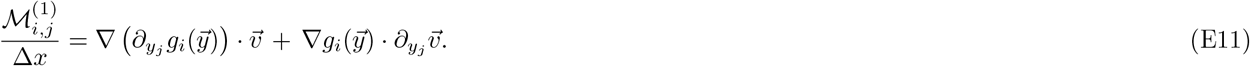

### E.1. Expansion of interaction matrix in environmental-organism models

Environmental-organisms models are a generalization of consumer-resource models proposed in ref.^36^ to capture a broader range of interactions between species and environmental factors. In particular, the model is defined as follows

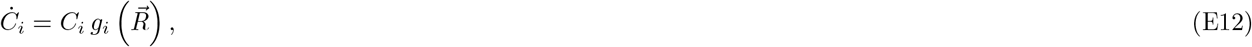

and

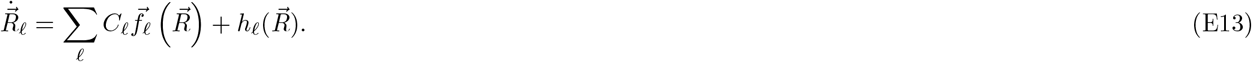

Here, the vector 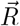 represents environmental traits that influence species growth, including resource availability, temperature, light, and other abiotic factors^36^.

Notably, this model does not include direct interactions between species populations (*C*_*i*_). As a result, the leading-order term in the time-expansion of the interaction measure vanishes:

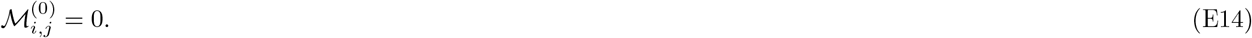

Thus, the first non-zero contribution is the next-order term 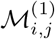, which captures fast, indirect interactions mediated by the environment. The expression for 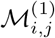 in Eq. (E11) simplifies when both indices refer to species since:

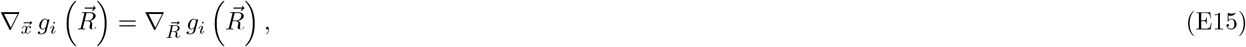

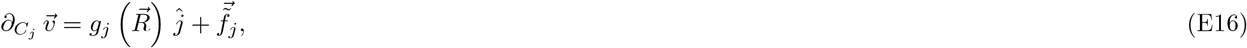

and

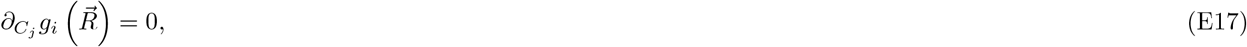

where 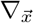 and 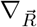 represent the nabla operator in the space of species and environmental vector jointly and the nabla operator in the space of environmental vector respectively. We also defined 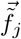 as a vector that has zeros in the species components and is equal to 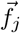 in the environmental components.

Substituting these expressions into Eq. (E11), we obtain the environmental-mediated interaction:

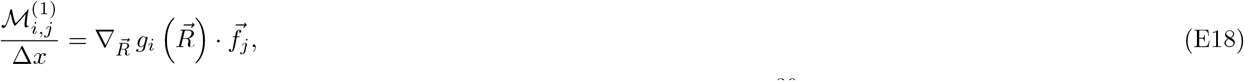

which corresponds to the instantaneous interaction matrix defined in Ref.^36^.

### E.2 Expansion of interaction matrix in consumer-resource models

In this section, we give explicit form for the expansion of the interaction matrix in the consumer-resource model defined in Eqs. (9) and (10). In particular, we focus in the interaction between two consumers. The gradient vector associated to consumers read

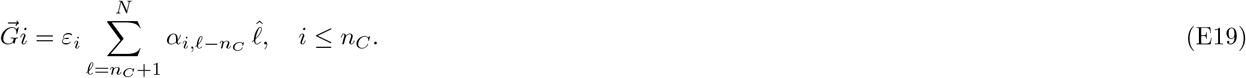

The coefficients *ℳ* ^(0)^, *ℳ* ^(1)^, and *ℳ* ^(2)^ are obtained inserting Eqs. (E7) and (E19) in Eq. (5):

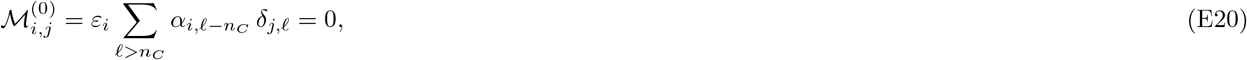

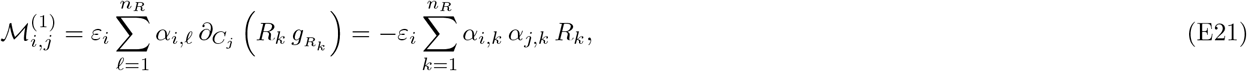

and

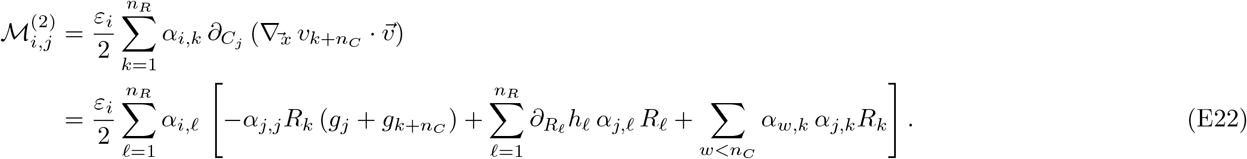

## Appendix F: Model inference from interaction matrix

If we were to measure the empirical interaction matrix, 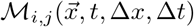, based on synthetic paths generated by the general model in Eq.(3) for small values of Δ*t*, Δ*x*, and *t*, we could access the limit

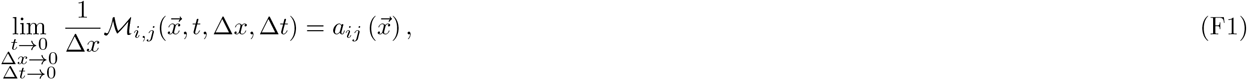

where

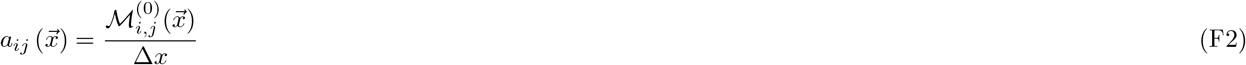

is the theoretical definition of the interaction matrix^30^. We can define the following vector function

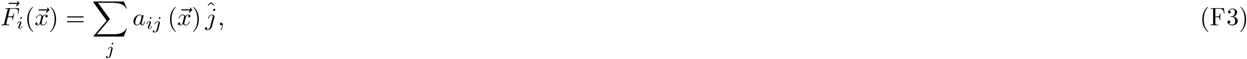

which, in virtue of Eq. (21) fulfills,

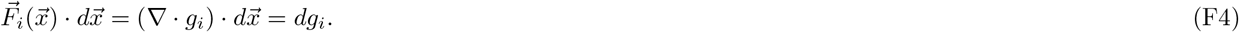

Therefore, the vector field 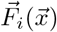 is conservative, i.e. 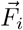 derives from a potential (scalar field) that we can identify as the per-capita growth rate of species *i*.

Now let us consider the situation in which we have real data with finite *t*, Δ*t*, and Δ*x*. From these data, we could compute the estimators

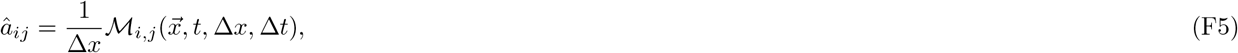

and

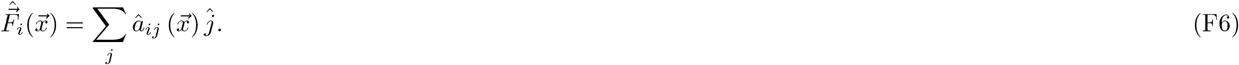

Then if 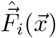 is conservative, i.e. if 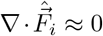, then Eq. (F4) can be seen as a differential equation for the per-capita growth rate with solution,

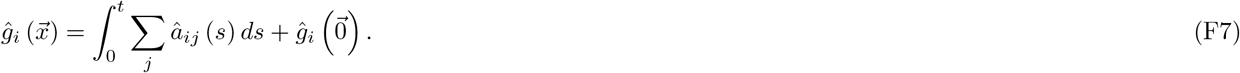

Moreover, considering again that 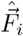 is conservative, then its integral does not depend on the integration path, and we can write

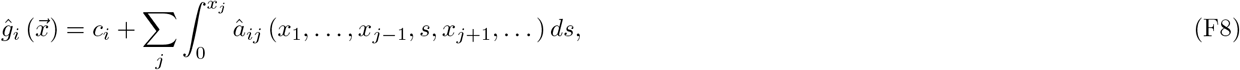

where 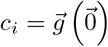 are integration constants, independent of the measure of interactions.

Using the definition of the local growth rate (Eq.(3)), the model inferred from the leading term of interactions would read

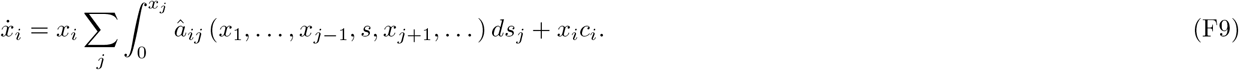

We note that the inference framework relies on the assumption that the empirical vector field, 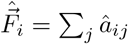, derives from a potential. However, it is not guaranteed that real data will satisfy this condition. In general, we would expect vector fields to have rotational contributions. According to the developed theory, this would imply that the species dynamics are not compatible with an equation of the form given by Eq. (3); or that the errors introduced by finite Δ*x*, Δ*t*, and *t* are too big to apply this method.

## Appendix G: Self-interactions in the geometric Brownian model

In this section, we compute self-interactions over paths generated with the geometric Brownian model,

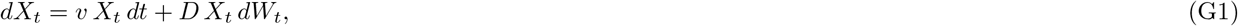

with solution

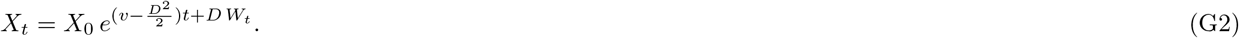

In this case, the instantaneous per-capita growth rate measured from a realization of the process with *X*_0_ = *x*_0_,

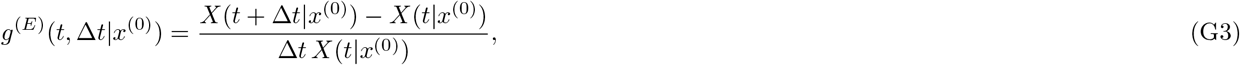

can be computed exactl using (G2),

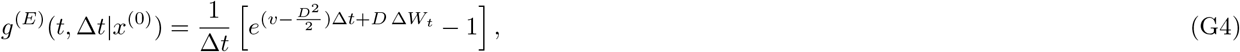

where

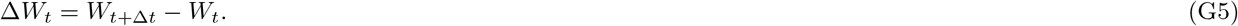

The empirical measure of interactions can also be evaluated from eq.(G4) as

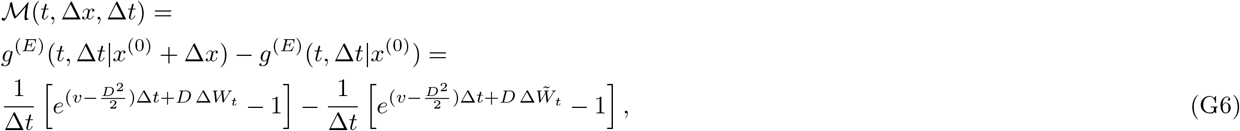

where *W*_*t*_ and 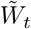 represent two distinct realizations of the Wiener process. For small Δ*t*, Eq. (G6) can be further simplified to

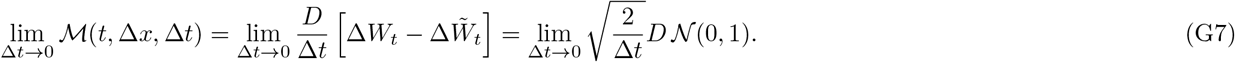

## Appendix H: Effect of noise correlations in growth rates estimates

In this section, we study per-capita growth rate estimates measured over trajectories generated by a process with colored noise:

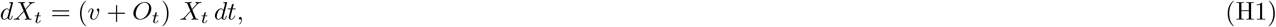

where *O*_*t*_ is an Ornstein-Uhlenbeck process,

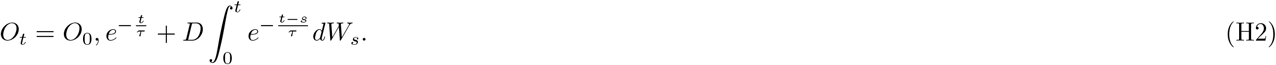

The process in Eq.(H1) has the solution

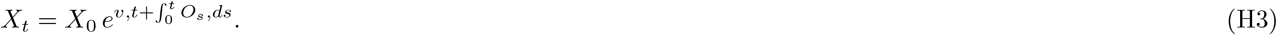

Inserting this solution into Eq.(1), we obtain the empirical growth rate over stochastic paths:

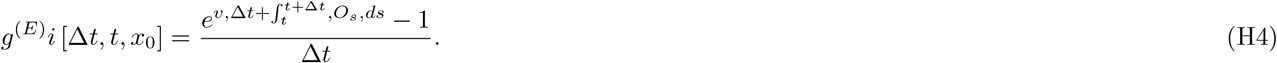

The limit Δ*t →* 0 can be safely computed,

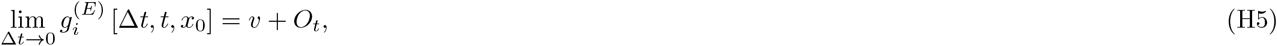

which is a well-defined random quantity corresponding to the deterministic growth rate with Gaussian perturbation.

If instead we generate data using uncorrelated noise,

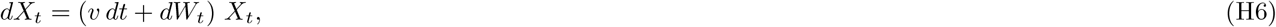

we obtain

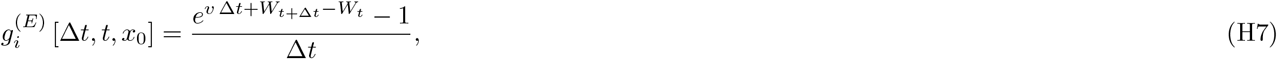

which has no well-defined limit as Δ*t →* 0, since

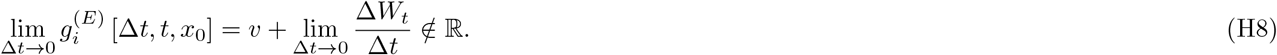

